# An Information Compass for Probabilistic Reasoning in the Brain

**DOI:** 10.64898/2026.10.07.757408

**Authors:** Juvenal Bosulu, Vijay Balasubramanian, Zachary P. Kilpatrick, Krešimir Josić, Joshua I. Gold

## Abstract

Probabilistic reasoning in the brain has been studied primarily using simple two-choice tasks that are experimentally and theoretically tractable but have unclear relevance to more complex problems. Here we generalize principles from these studies using information geometry, which treats probability distributions (“beliefs”) as points on statistical manifolds whose geometry captures relationships between those beliefs. We introduce an “information compass,” a context-dependent, one-dimensional axis along which an arbitrarily complex belief is reinforced or repudiated. Projecting high-dimensional belief updates onto this axis casts even complex decisions into familiar forms of deliberation and commitment used for simpler decisions. Our framework provides a unified account of how biological and artificial systems can use geometric forms of probabilistic reasoning to make decisions across a range of tasks and domains.

## Introduction

Higher brain function depends on the ability to form, update, and use beliefs about the world. These beliefs can be represented as probability distributions over possible external or internal states. For example, perceptual inference uses sensory evidence to form and update beliefs that inform judgments about what we see, hear, feel, taste, and smell [1, 2]. Likewise, active inference transforms uncertain beliefs about possible actions and their consequences into committed behavioral responses [3]. In general, these forms of probabilistic reasoning require a balance of: 1) deliberation, in which beliefs evolve in response to new information, contexts, or goals; and 2) commitment, in which those beliefs are used to inform judgments and actions.

Our understanding of how deliberation and commitment are implemented in the brain is based largely on studies using decisions between two alternatives [4]. This design reduces the decision process to a tractable one-dimensional problem. For example, binary beliefs can be represented by a single scalar decision variable, such as the log-odds ratio log [*p/*(1 − *p*) ], where *p* is the probability assigned to one alternative [5, 6]. Deliberation corresponds to movement of this decision variable along a single axis, toward one alternative and away from the other. Commitment is often assumed to occur when the decision variable reaches a bound, which is a point on this axis. This formulation underlies many normative and mechanistic models of decision-making and has close ties to both behavioral and neural findings [7–9].

However, extending these principles to more complex, real-world beliefs remains challenging, particularly when probability distributions vary widely in dimensionality, structure, and interpretation. For example, a friend’s grimace after tasting your cooking may update a simple belief you have about whether they liked it, while leaving ambiguous a richer belief about what went wrong (too salty? too spicy? undercooked?). More generally, high-dimensional belief spaces can obscure the context-relevant directions important for deliberation and commitment, which complicates both the brain’s ability to implement those processes and researchers’ ability to understand them.

Here we propose that many principles of deliberation and commitment, including those derived from studies of simple, two-alternative decisions, can be cast in information-geometric terms [10–12] that generalize naturally to more complex problems. We formalize this generalization with an “information compass” that measures, for any belief, whether, and by how much, a given update reinforces or repudiates that belief. This approach thus recasts high-dimensional (multi-alternative) belief updates into familiar, two-alternative forms. We then illustrate the usefulness of this approach with applications spanning decision-making, goal-seeking behaviors, learning, and information sampling.

## Results

### The information compass

A foundational idea in information geometry is that any probability distribution, *P*, can be treated as a point on a statistical manifold that represents the space of possible beliefs within a given probabilistic model (here, discrete distributions over a finite set of alternatives; extensions to other exponential families are noted in the Discussion). Thus, updating a belief corresponds to moving from one point on this manifold to another. Information geometry provides a framework for characterizing these movements, allowing belief updates to be described in familiar geometric terms like distance and direction (Methods M1).

The direction of a belief update governs its effect, including whether it reinforces or repudiates the current belief. This distinction is not apparent using common measures of belief updating, such as the Kullback–Leibler (KL) divergence or Bayesian surprise, because they are unsigned and thus report only how much a belief changed, not in which direction [13]. To quantify this directional component, we introduce the “information compass” (Fig. 1). A magnetic compass uses north as a reference to navigate directions in the real world, including along lines of longitude that are the straightest paths from the current location to either pole. Likewise, an information compass uses a point on a statistical manifold as a reference to assess and guide the direction of belief updating, including along the straightest path from the current belief to that reference belief.

**Fig. 1.**
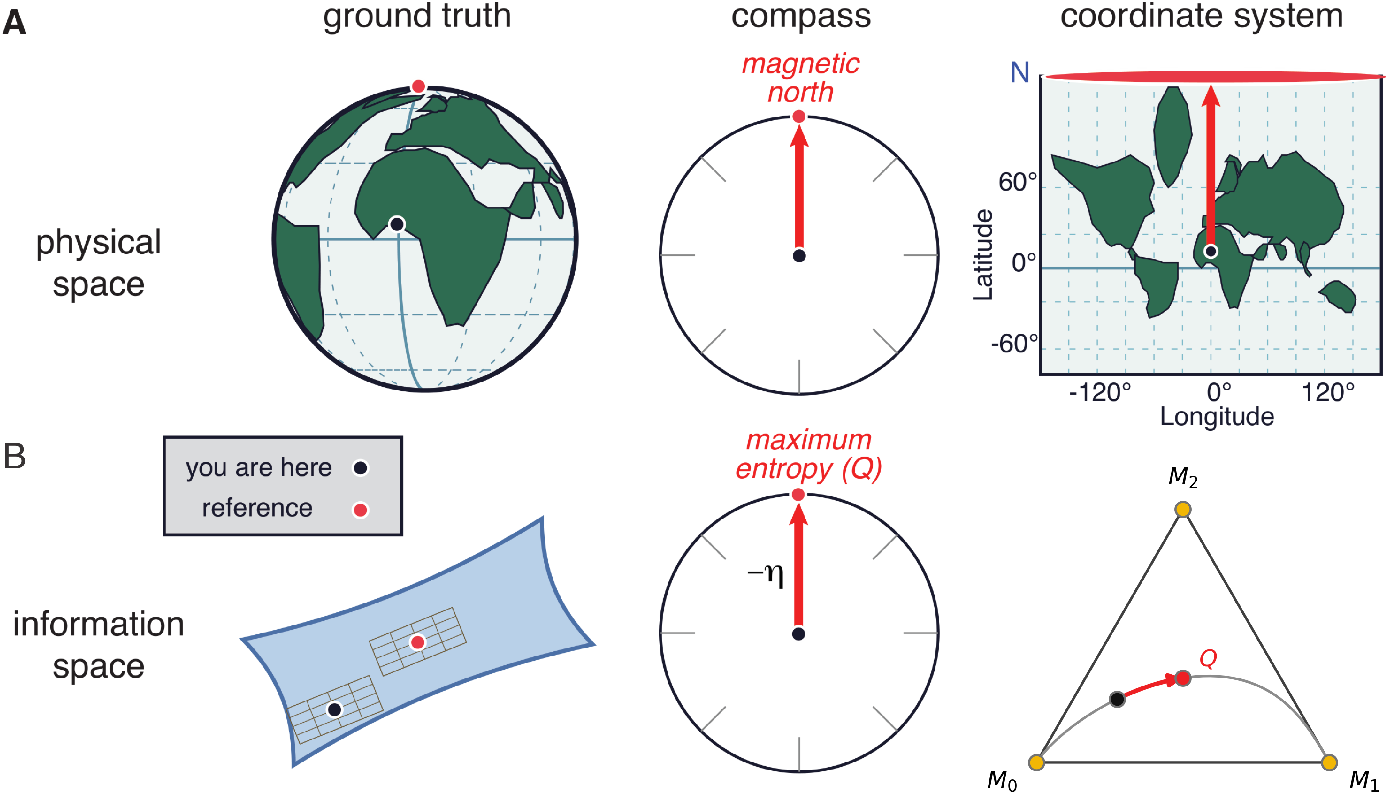
Comparison of a magnetic (A) and information (B) compass. **A.** A location on the Earth’s surface is a point on a spherical manifold. A magnetic compass at that point orients towards magnetic north. Navigating with the compass involves aligning it with a map expressed using a particular coordinate system (e.g., a flat map that stretches distances near the poles). **B**. A belief is a point on a statistical manifold, where distances and angles are measured using a metric (typically the Fisher information metric) that defines “closeness” in terms of how distinguishable two distributions are. Each point has a tangent space whose basis vectors define the local coordinate directions (the grids in the figure). The information compass at a belief *P* is the tangent vector pointing toward the maximum-entropy belief (*Q*), which equals the negative of *P* ‘s *e*-(natural-parameter) coordinates. It lies along a geodesic (a straight line in *e*-coordinates) that runs from the maximum-certainty belief favored by *P* (yellow circle, *M*_0_), through *Q*, to an alternative maximum-certainty belief (here *M*_1_). This geodesic can appear curved in *m*-coordinates (the simplex, shown for three-state beliefs on the right; Methods M1).

Making this idea precise requires specifying what counts as a straight path on a statistical manifold, which in turn depends on how beliefs are described. Consider beliefs about *n* possible states, *s*_0_, …, *s*_*n−*1_. Information geometry uses two different coordinate systems to represent these beliefs, each suited to describing a different kind of belief change. In exponential (*e*) coordinates, a belief is described by its log-odds relative to a reference state, *η*_*i*_ = ln[*P* (*s*_*i*_)*/P* (*s*_*n−*1_)], *i* = 0, …, *n* − 2, which generalizes the log-odds decision variable used for two-alternative tasks. In these coordinates, Bayesian updating about a fixed state of the world is simple addition: each new piece of evidence adds its log-likelihood ratio to the current log-odds. In mixture (*m*) coordinates, a belief is described directly by its probabilities, *p*_*i*_ = *P* (*s*_*i*_), *i* = 0, …, *n* − 2 (*p*_*n−*1_ is determined because probabilities sum to one), which places it in the familiar probability simplex (Extended Data Fig. 1). In these coordinates, hedging between possibilities is simple averaging: if the world remains in the state described by *P*_1_ with probability 1 − *h* but has switched to the state described by *P*_2_ with probability *h*, the appropriate belief is (1 − *h*)*p*_1_ + *h p*_2_. Each coordinate system thus has its own notion of a straight path, or geodesic, through belief space (*e*-geodesics for evidence accumulation, *m*-geodesics for mixing), and a path that is straight in one generally appears curved in the other (Fig. 1B).

The information compass is based on the *e*-geodesic that passes through an “origin” belief (*P*_*o*_) and the reference belief. Here we use a reference belief corresponding to complete uncertainty, or more precisely the “maximum-entropy” belief (which we refer to as *Q*, expressed as *p*_*i*_ = 1*/n* in *m*-coordinates and *η* = 0 in *e*-coordinates; other reference beliefs are considered below). Because *Q* lies at the origin of *e*-coordinates, this geodesic consists of all beliefs whose *e*-coordinates are scaled versions of those of *P*_*o*_, *η* = *t η*_*o*_. Every point on this line thus has a simple interpretation in terms of evidence. Values of *t >* 1 correspond to beliefs that favor the same states as *P*_*o*_ but with greater confidence, as if more of the evidence that produced *P*_*o*_ had been accumulated. Values of 0 *< t <* 1 correspond to weaker versions of *P*_*o*_, reaching total uncertainty (*Q*) at *t* = 0. Negative values correspond to beliefs whose evidence runs counter to that of *P*_*o*_, including the “opposite” belief 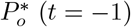, whose evidence exactly cancels that of *P*_*o*_ (Methods M3). At its extremes, the line ends at certainty in the states that *P*_*o*_ considers most likely (*t*→ ∞ ) and least likely (*t*→ −∞). These endpoints are vertices of the simplex unless *P*_*o*_ assigns equal probability to more than one most- or least-likely state (Fig. 1B; Methods M1).

For two-alternative beliefs, this *e*-geodesic is the entire (one-dimensional) belief space. For more alternatives, it is a one-dimensional subspace, and a general update from an origin belief, *P*_*o*_, to a destination belief, *P*_*d*_, need not lie along it. Rather, in *m*-coordinates the update is a displacement, *p*_*d*_ − *p*_*o*_, that can point in any direction within the probability simplex. We can quantify its component along the compass axis, which we refer to as the information with respect to the compass, *i*_*c*_, as:

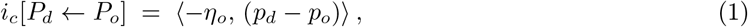

where ⟨ ·, · ⟩ denotes the ordinary dot product. In general, measuring the angle between two directions on a statistical manifold requires a metric (the Fisher information metric) that varies from point to point. However, the ordinary dot product between one displacement expressed in *e*-coordinates and the other displacement expressed in *m*-coordinates, as in Eq. 1, equals the inner product defined by that metric [11] (Methods M1). Equivalently, *i*_*c*_ can be computed from probabilities alone, as the change in the probability assigned to each state, weighted by how surprising that state was under the origin belief (− ln *P*_*o*_(*s*_*i*_); Methods M2, Eq. 14). Thus, *i*_*c*_ is positive when an update shifts probability toward states that the origin belief found more surprising than average, and negative when it shifts probability toward states that it found less surprising than average.

For small updates, *i*_*c*_ is the projection of the update (*p*_*d*_ − *p*_*o*_) onto the compass heading (−*η*_*o*_), measured in the natural geometry of the manifold. For updates of any size, *i*_*c*_ retains an exact informationtheoretic form (Methods M2), equaling the KL divergence from *P*_*o*_ to *P*_*d*_ minus the gain in certainty (the reduction in entropy, or increase in negentropy, Δ*φ*) from *P*_*o*_ to *P*_*d*_:

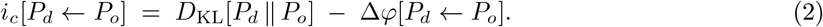

The KL divergence measures how much information distinguishes the destination from the origin belief and is always non-negative. In contrast, Δ*φ* is signed, with positive and negative values corresponding to increases and decreases in certainty, respectively. Thus, unlike unsigned measures of surprise, *i*_*c*_ captures how much of an update moves along the compass axis in either direction, beyond any change in overall certainty (Fig. 2A,B; Extended Data Fig. 3).

**Fig. 2.**
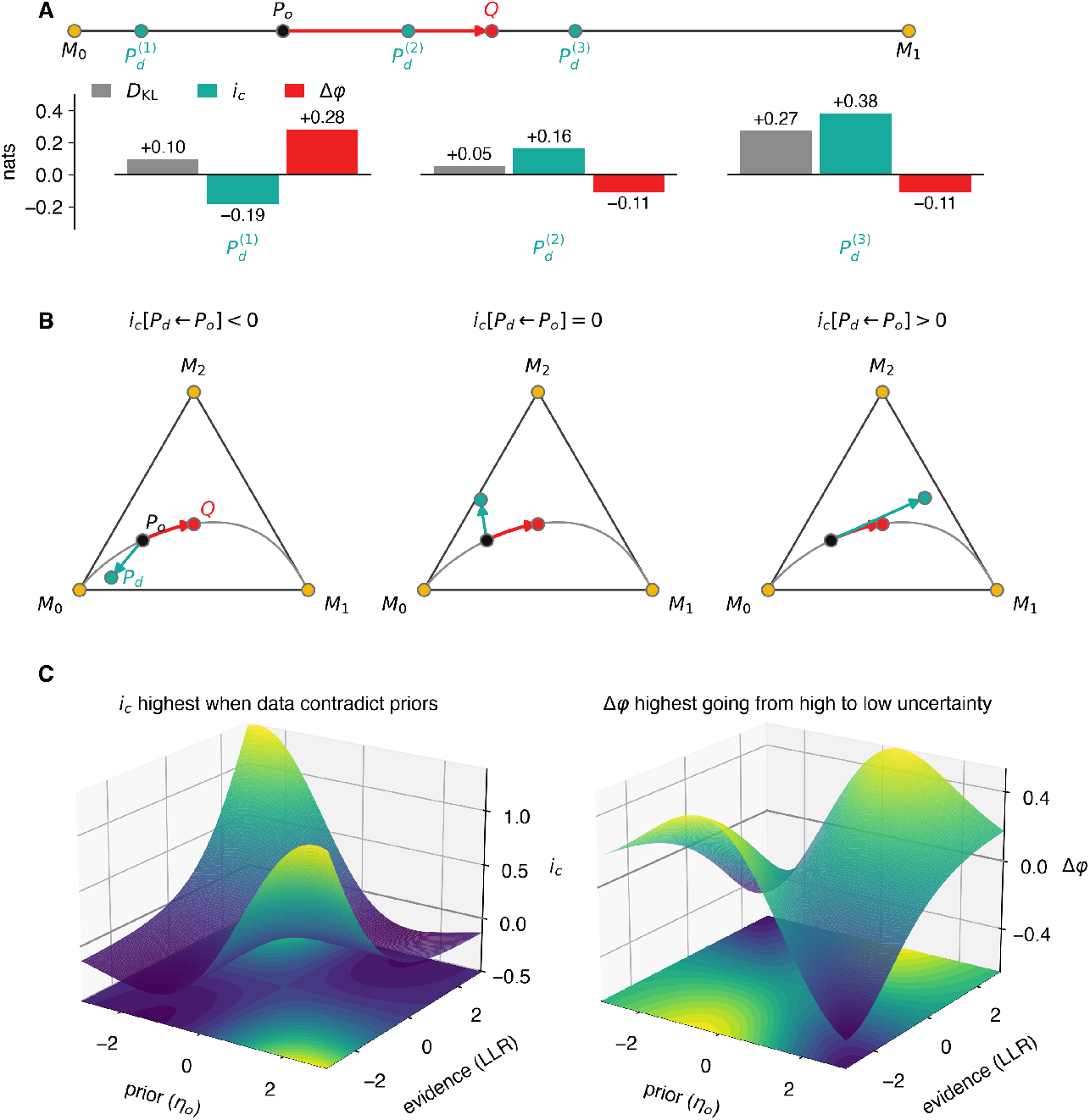
Information with respect to the compass (*i*_*c*_). **A.** The geometry of two-state beliefs shown on a 1D simplex. *M*_0_ and *M*_1_ are the two minimum-entropy beliefs. *Q* is the maximum-entropy belief. *P*_*o*_ is an origin belief that favors *M*_0_ (*P*_*o*_(*s*_0_) = 0.75). Bars show, for three destination beliefs (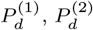, and 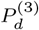), the KL divergence from *P*_*o*_ (gray) and its two components (Eq. 2), *i*_*c*_ (teal) and Δ*φ* (red). 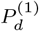 moves toward *M*_0_ and reinforces *P*_*o*_ (*i*_*c*_ *<* 0). 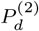 moves toward *Q*, and 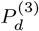 crosses it to favor *M*_1_, both repudiating *P*_*o*_ (*i*_*c*_ *>* 0) while losing certainty (Δ*φ <* 0). **B**. The geometry of three-state beliefs shown on 2D simplexes, for three different values of *i*_*c*_, as indicated. **C**. *i*_*c*_ (left) and Δ*φ* (right) for twostate Bayesian updating, in which a prior (origin) belief *P*_*o*_ becomes a posterior (destination) belief *P*_*d*_ by adding evidence in *e*-coordinates, plotted as a function of the prior’s natural parameter (*η*_*o*_) and the evidence log-likelihood ratio (LLR). The shaded projection on the floor of each panel repeats the same surface viewed from directly overhead. In general, *i*_*c*_ is high and positive when the evidence contradicts the prior and becomes negative when the evidence reinforces the prior. Δ*φ* is the entropy reduction across the update, so it is largest when an uncertain prior (*η*_*o*_ ≈ 0) is resolved by strong evidence in either direction, and most negative when evidence pushes a confident prior back to near-maximal uncertainty (LLR ≈ −*η*_*o*_). Evidence strong enough to overshoot *Q* and confidently reverse the belief raises Δ*φ* again.

The sign of *i*_*c*_ represents the direction of the update and thus a logical interpretation of its effect on the origin belief. Negative values, which move away from *Q* and towards the nearest maximum-certainty belief, represent reinforcement of *P*_*o*_ (Fig. 2C). This kind of belief strengthening occurs when, for example, evidence is accumulated to achieve sufficient confidence to commit to a decision [8, 14– 19]. In contrast, positive values, which move toward *Q* (where all structure in the belief is lost) and potentially beyond it toward states that *P*_*o*_ considered unlikely, represent repudiation of *P*_*o*_. This kind of belief update underlies changes of mind and many forms of learning that involve violations of current beliefs or expectations (e.g., prediction-error learning; 20–29).

We next apply the information compass to two key components of belief updating: 1) deliberation, which can be thought of as a path over a statistical manifold of beliefs; and 2) commitment, which can be thought of as the location(s) on the manifold where deliberation stops.

### Deliberation as a path in belief space

Deliberation between two alternatives corresponds to movements along a single dimension that can be readily interpreted in terms of changing support both for and against each alternative [30]. In contrast, adding even one more alternative creates a higher-dimensional belief manifold in which belief updates have multiple degrees of freedom, making them potentially more difficult to compute, interpret, and control [31]. Here we use two examples to illustrate how the information compass can establish task-relevant, one-dimensional axes that restore the simplifying logic and symmetries of two-alternative tasks in higher-dimensional settings [32].

#### Evidence accumulation in changing environments

Effective decision-making in changing environments requires maintaining and updating beliefs about the nature of the changes. For binary choice tasks in which the changes correspond to a fixed rate of switching between two states (at a “hazard rate” *h*), optimal, accuracy-maximizing decisions depend on ongoing, hazard-dependent updates of beliefs about the current state [33, 34]. Specifically, under these conditions the Bayesian belief after observing the *i*-th piece of evidence (*θ*_*i*_, representing the logarithm of the posterior odds of one alternative versus the other) is updated from the previous timestep as:

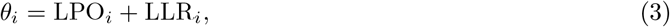

where

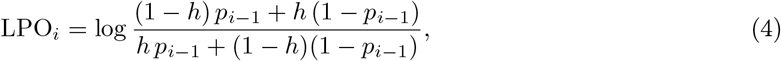

is the log-prior-odds, expressed in terms of the posterior probability of a particular alternative at the previous step (*p*_*i−*1_), after taking into account the possibility that the state changed (with probability *h*) since that step; and LLR_*i*_ is the log-likelihood ratio of the new evidence.

This update generalizes directly to beliefs over any number of states *n*, where on each timestep the current true state can switch to any one of the *n* − 1 other states, with total probability *h* distributed uniformly across those states (a special case of a more general model that allows a separate switching rate for every ordered pair of states [34]). This generalization uses the same two main components of the *n* = 2 derivation, with clear mappings onto the dual coordinate systems of information geometry.

The first component is a mixture over all possible previous states, weighted by the hazard rate *h*. This mixture is straightforward to compute in *m*-coordinates, as a convex combination that produces the new prior, *P*_*h*_:

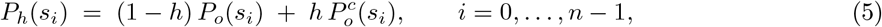

where 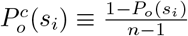 is the belief that would result if a switch were certain (*h* = 1): each state receives an equal share of the probability that *P*_*o*_ assigned to the other states. *P*_*h*_ thus interpolates between *P*_*o*_ (*h* = 0) and 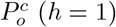, before any new evidence is considered.

The second component then combines this new prior with the new evidence. This combination is straightforward to compute in *e*-coordinates, as a vector addition that produces the new posterior *η*_*d*_:

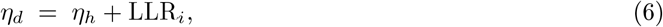

where *η*_*h*_ denotes the *e*-coordinates of *P*_*h*_ from Eq. 5 (Methods M1; Fig. 3A). At *n* = 2, 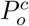 is simply the other state, so Eq. 5 reduces exactly, term for term, to Eq. 4 (Methods M3).

**Fig. 3.**
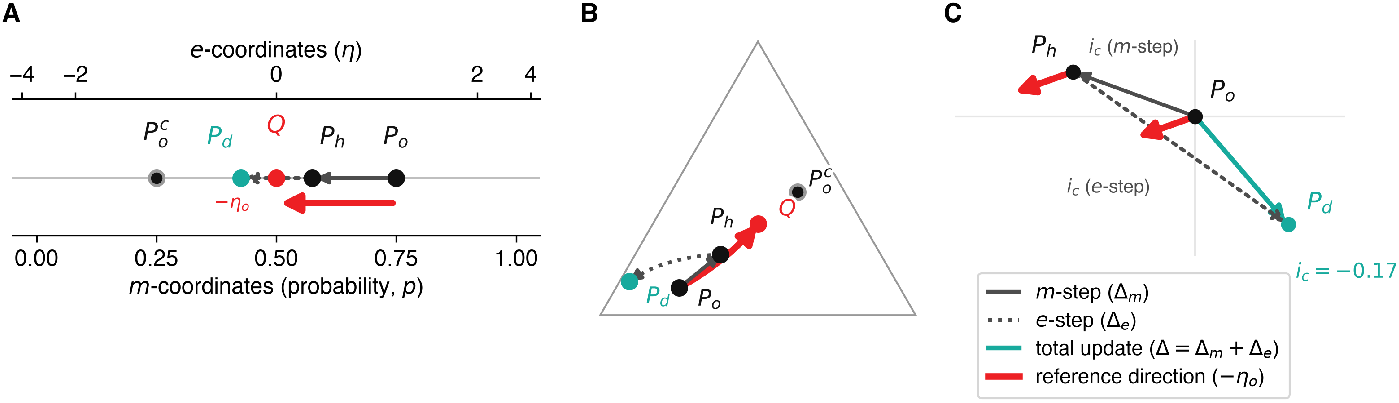
Using the information compass to track normative, adaptive prior updating in changing environments. **A.** For two-state beliefs (*n* = 2), the hazard-dependent *m*-step (thin solid, *P*_*o*_ to *P*_*h*_; Eq. 5) and the evidence-dependent *e*-step (thin dotted, *P*_*h*_ to *P*_*d*_; Eq. 6) are consecutive segments of a single axis, shown with *m*- and *e*-coordinate readouts below and above. 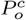 (the belief if a switch is certain), *Q*, and the reference direction −*η*_*o*_ (thick red, on its own row below) are marked for reference. **B**. For three-state beliefs (*n* = 3), shown on the simplex, the same two steps are no longer collinear, because the *m*-step (thin solid) is straight in *m*-coordinates, heading directly to *Q* (and, if continued to *h* = 1, to 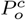), whereas the *e*-step (thin dotted) is straight in *e*-coordinates and so appears curved here. The reference direction (thick red) is the *e*-geodesic from *P*_*o*_ to *Q*. **C**. The same *n* = 3 update in reduced *m*-coordinates (the two axes represent the probabilities of the first two states, with *P*_*o*_ at the origin), so the *m*-step displacement (Δ_*m*_, thin solid) and the *e*-step displacement (Δ_*e*_, thin dotted) appear as vectors in the plane, summing to the full update Δ (teal). Each of Δ_*m*_ and Δ_*e*_ is projected onto the reference direction −*η*_*o*_ (thick red, redrawn from each tail), and the labeled *i*_*c*_ (*m*-step) and *i*_*c*_ (*e*-step) projections sum to the *i*_*c*_ of the full update (*i*_*c*_ = −0.17 here; Eq. 7; Methods M3).

This use of dual coordinates eases the computational burden of extending a two-alternative update to many alternatives. Because the mixing step (Eq. 5) is linear in *m*-coordinates and the evidence step (Eq. 6) is additive in *e*-coordinates, neither step needs to be re-derived as *n* grows; only the vectors get longer. In contrast, a previous formulation that uses only log-probability space requires additional computations as *n* grows, including new coupling terms and log-likelihood ratios computed for each pair of alternatives [34].

However, these computational benefits raise another challenge. If the two steps operate in different coordinate systems, for *n >* 2 it may not be clear if and how each step contributes to the overall update, or how far the overall update progresses towards or away from maximum uncertainty (*Q*; Fig. 3B). These questions can be answered using the information compass. Because *i*_*c*_[*P*_*d*_ ← *P*_*o*_] = (−*η*_*o*_) · (*p*_*d*_ − *p*_*o*_) (Eq. 1) is linear in the destination belief’s *m*-coordinates for a fixed origin *P*_*o*_, it can be applied separately to the *m*-step and the *e*-step, using *P*_*o*_’s own reference direction (−*η*_*o*_) throughout rather than recomputing it at *P*_*h*_. The two contributions then add exactly to the *i*_*c*_ of the full update (Fig. 3C; Methods M3):

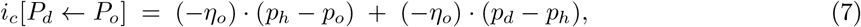

where *P*_*d*_ is the posterior distribution obtained after the full update (*η*_*d*_ in *e*-coordinates).

In short, *i*_*c*_ turns a multi-step update across two different coordinate systems into a single running tally on one fixed axis (here defined by the belief before each update; Methods M3 describes an alternative that keeps one reference belief fixed across all updates). This tally can be used to compare the contributions of the two steps in terms of their net effect on the belief. The hazard-dependent *m*-step, which reflects expected volatility, always has *i*_*c*_ ≥ 0 as long as switching does not favor particular states (as for the uniform switching considered here): without new evidence, the possibility that the world has changed can only weaken the current belief, never reinforce it (Methods M3). In contrast, the evidence-dependent *e*-step can have either sign. The compass thus splits any change of mind into a part attributable to expected volatility and a part attributable to new evidence, for any number of alternatives. This split makes it possible to ask, for example, whether a particular change of mind was driven mainly by the expectation that the world might have changed or by evidence that it had.

#### Context-dependent motivated behaviors

Real-world belief updates must often account for not just multiple alternatives but also relationships among multiple variables whose joint distribution defines context-dependent goals. Under these conditions, the information compass can be realigned toward the current goal. For any reference belief *P*_*r*_ (e.g., a goal to be approached or avoided), the difference between the reference and origin beliefs in *e*-coordinates, Δ_*r*_*η*_*o*_ = *η*_*r*_ − *η*_*o*_, defines a compass toward *P*_*r*_ (Methods M2):

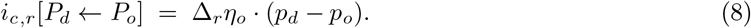

This quantity measures the extent to which an update from *P*_*o*_ to *P*_*d*_ moves toward *P*_*r*_, collapsing a potentially high-dimensional, multi-goal update onto a single axis.

We illustrate the usefulness of this formulation in terms of relationships between physiological need and feeding behaviors [35–37]. Rats normally avoid cues paired with highly concentrated salt and show strong disgust-like reactions to it [37]. However, in a novel sodium-deficient state, the same rats rapidly seek out those same cues, despite never having experienced their positive hedonic value under deficiency [38–41]. Thus, need can produce instantaneous motivational value independent of learned experience, which is thought to reflect a form of active inference that prioritizes cues capable of fulfilling prior physiological preferences [42, 43].

We model this phenomenon using two beliefs, each with its own preferred state: sodium state *S* = {*s*_0_, *s*_1_}, where satiation (*s*_1_) is preferred, and discomfort *D* = {*d*_0_, *d*_1_}, where its absence (*d*_0_) is preferred (Fig. 4A). Together, these preferences define a joint goal, *P*_*r*_ ≈ *P* (*s*_1_, *d*_0_). We assume that the only available action, consuming the concentrated salt, always produces a belief that is only partly preferred, *P*_*d*_ ≈ *P* (*s*_1_, *d*_1_) (sated but uncomfortable). The value of this action, measured as *i*_*c*_ with respect to the goal (Eq. 8), depends strongly on the origin belief (Fig. 4B,C). When the origin is highly deprived, *i*_*c*_ is large and positive, reflecting a strong pull toward the salt despite the discomfort it brings. As deprivation resolves, *i*_*c*_ does not simply shrink toward zero but changes sign, becoming negative near satiation.

**Fig. 4.**
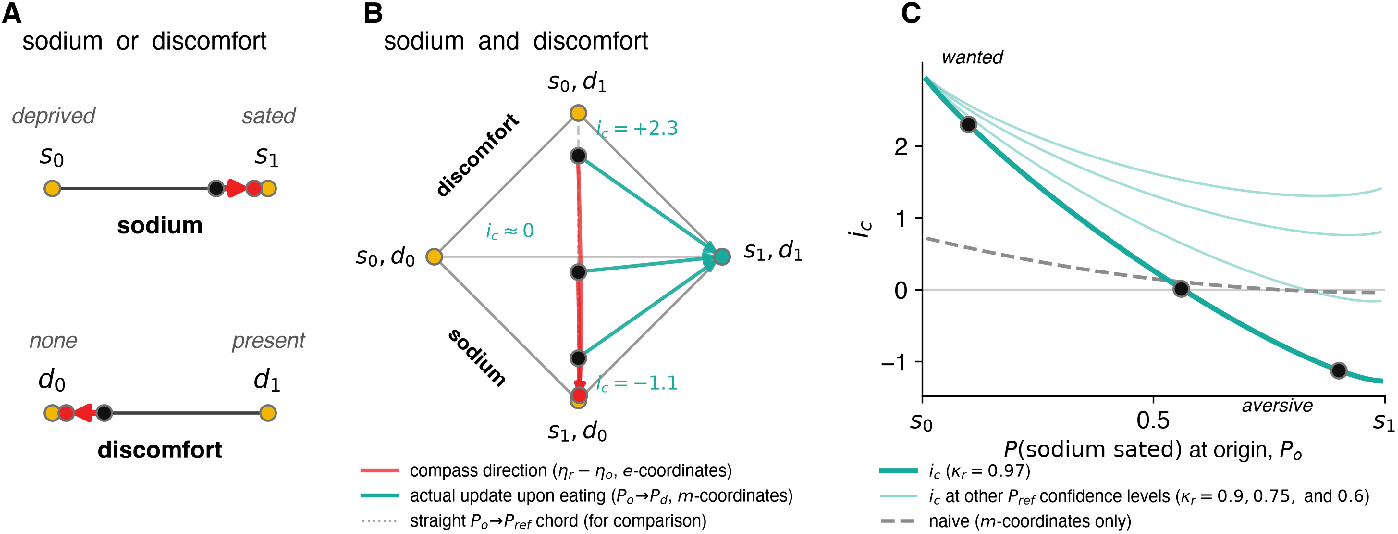
Using the information compass to guide context-dependent, goal-directed behaviors. **A.** 1D simplexes for sodium state *S* = {*s*_0_, *s*_1_} and discomfort *D* = {*d*_0_, *d*_1_}, each with a compass from an origin belief (black) realigned to its preferred state (*s*_1_, sated; *d*_0_, no discomfort). **B**. 3D simplex (a tetrahedron drawn in 2D) for the joint distribution over *S* and *D*. Origin beliefs (black dots) lie on the axis from deprivation with discomfort, *P* (*s*_0_, *d*_1_), to the goal *P*_*r*_ ≈ *P* (*s*_1_, *d*_0_). Consuming salt moves the belief toward *P*_*d*_ ≈ *P* (*s*_1_, *d*_1_) (teal arrows, with *i*_*c*_ values indicated). Red arrows: each origin’s compass toward the goal (*η*_*r*_ −*η*_*o*_, curved in the simplex); gray dotted line: the straight *P*_*o*_ →*P*_*r*_ chord. Beliefs are held with confidence *κ*_*r*_ = 0.97 (goal) and *κ*_*o*_ = 0.88 (origins) rather than certainty, because a compass pointing at a vertex would be infinite. **C**. *i*_*c*_ (teal) as a function of the origin belief, *P* (sodium sated); black dots mark the origins in B. Gray dashed line: the same comparison made directly on probabilities (*m*-coordinates only). Both change sign as the origin moves toward satiation, but at different points, and only *i*_*c*_ captures the outsized pull of acute need. Faint curves: *κ*_*r*_ = 0.90, 0.75, and 0.60; the aversive switch requires *κ*_*r*_ *> κ*_*o*_.

Thus, the same action, with the same outcome, is strongly favored under deprivation but actively disfavored once sodium need is met. This reversal occurs when the goal considers the salt’s outcome (sated with discomfort) less likely than a sated origin does, which here corresponds to holding the goal with greater confidence than the origin (*κ*_*r*_ *> κ*_*o*_; Fig. 4C). Otherwise, a sated origin is already more certain of the preferred state than the goal itself, so the compass points back toward less certainty and consuming the salt remains favored.

This sensitivity to context, previously explained either as need instantly making a cue “wanted” (incentive salience; 41) or as need acting as a surprise signal that pulls behavior toward preferred states [43], arises here because *i*_*c*_ includes the term *D*_KL_[*P*_*d*_ ∥*P*_*o*_], which grows sharply as *P*_*o*_ approaches a nearcertain belief that *P*_*d*_ moves away from (here, extreme sodium depletion). This outsized sensitivity to acute need is far weaker in an otherwise identical quantity computed entirely in *m*-coordinates (dashed line in Fig. 4C), which, unlike *i*_*c*_, depends on the arbitrary choice of coordinates. This example thus illustrates how the information compass can capture interactions among multiple goals.

### Commitment as locations in belief space

In general, belief updating in the brain is an ongoing process with no clear starting or ending point. However, along the way this process is used to achieve a multitude of goals, such as making accurate perceptual judgments or committing to appropriate actions. Here we show that the information compass can determine if and when such a goal has been reached by recasting commitment as a comparison between two beliefs (the current belief and a commitment, or bound, belief), regardless of belief complexity. We illustrate this approach for multi-alternative decisions and information-gathering problems.

#### Multi-alternative decision rules

Many prominent models of decision-making (e.g., the Drift-Diffusion Model, or DDM) terminate the decision process using a bound on accumulated evidence (the decision variable) that governs trade-offs between decision speed, emphasized at low bounds, and accuracy, emphasized at high bounds [7–9]. For two-alternative decisions, two bounds are used, corresponding to the minimum confidence needed to commit to each alternative (e.g., “I will decide A when I believe A strongly enough”). Because the manifold is one-dimensional, each bound is simply a point, and thus a belief, that the decision variable crosses to commit.

For decisions involving more than two alternatives, a bound on accumulated evidence is not a point (i.e., not a single belief) but a higher-dimensional manifold of multiple beliefs. Such a manifold can, in principle, be derived from a specific speed-accuracy (or other) objective, but doing so generally requires solving a value function whose complexity grows with the number of alternatives [31, 44]. Classic multialternative sequential-sampling models avoid this complexity, and may not always follow normative principles, by instead using independent, fixed thresholds on each alternative separately (e.g., a race between accumulators, or the multi-hypothesis sequential probability ratio test) [45, 46], which treats the bound as a set of scalar criteria rather than as a single belief.

Here we propose an approach that keeps the bound as a single belief regardless of the number of alternatives. The bound is an origin belief (*P*_*o*_), the decision variable is a destination belief (*P*_*d*_), and the bound is crossed when *i*_*c*_[*P*_*d*_ ← *P*_*o*_] becomes negative, such that *P*_*d*_ “confirms” *P*_*o*_. As in the two-alternative case, commitment thus reduces to a comparison against a single belief. The compass of *P*_*o*_ defines the committed region (a half-space in *m*-coordinates) as all beliefs that represent equal or greater confidence in the state(s) favored by *P*_*o*_ (Fig. 5). When *P*_*o*_ treats the other alternatives symmetrically, as in Fig. 5, this region is simply a fixed threshold on the favored alternative’s probability, as in the multi-hypothesis sequential probability ratio test. The compass formulation extends this criterion to asymmetric bounds and, as described next, to bounds that change with urgency.

**Fig. 5.**
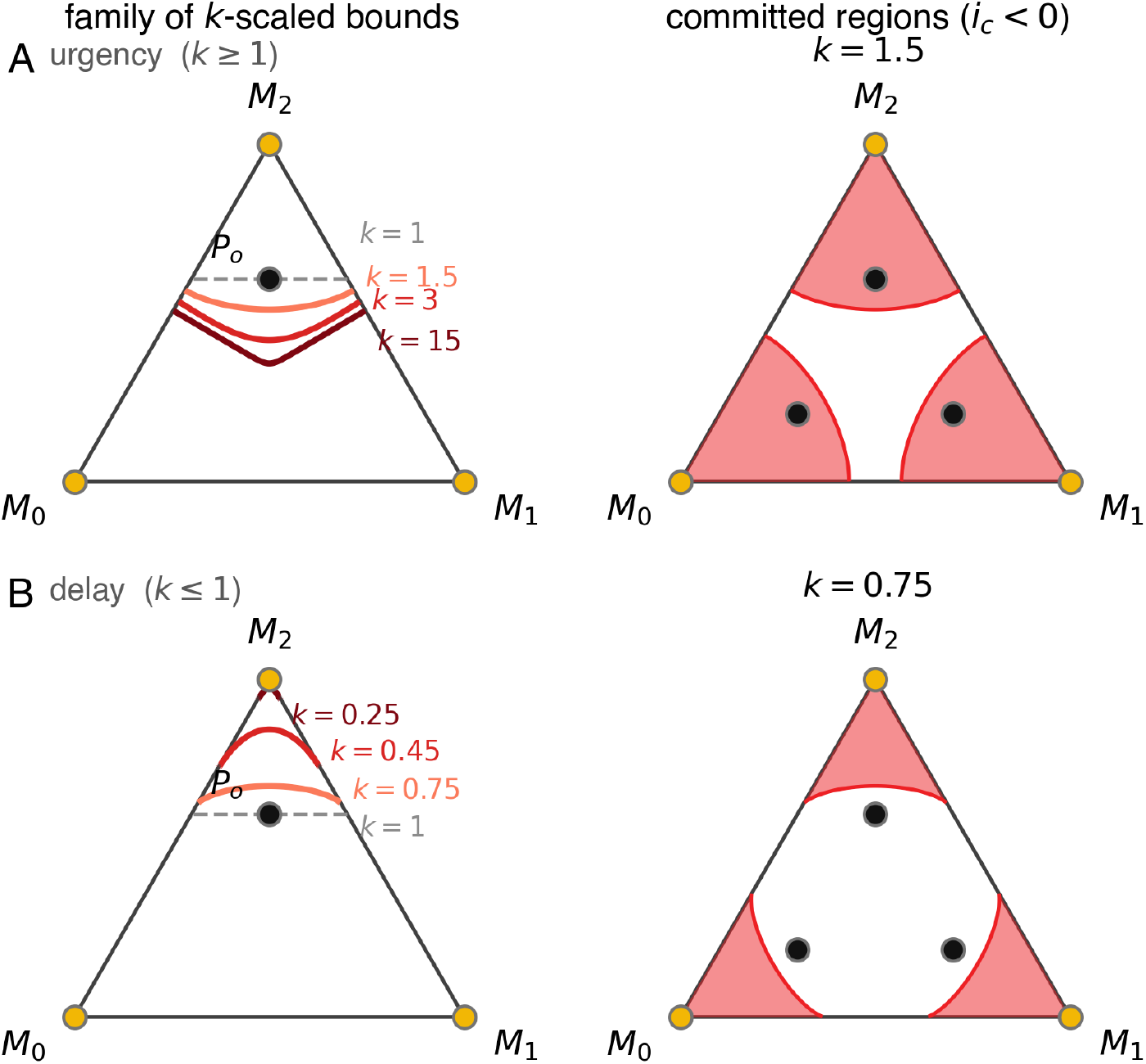
Using the information compass to define decision bounds for multi-alternative decisions. *Left* panels in both rows use the bound belief *P*_*o*_ = (0.2, 0.2, 0.6) (black dot), which favors vertex *M*_2_ symmetrically over *M*_0_ and *M*_1_. *Right* panels use three symmetric copies of *P*_*o*_ (black dots), one favoring each of *M*_0_, *M*_1_, and *M*_2_ in turn, so all three alternatives are governed by an equivalent bound. **A**. *Left:* Family of bounds for *k* ≥ 1 (urgency). Scaling the decision variable’s own compass by *k* = 1.5, 3, 15 (red curves, increasing saturation) pulls the *i*_*c*_=0 bound away from *P*_*o*_ and toward the maximum-uncertainty points reachable along *M*_2_’s two edges. The gray dashed line is *P*_*o*_’s own unscaled *k* = 1 bound. *Right:* Committed regions (shaded) for *k* = 1.5, one per alternative, each consisting of all beliefs *P*_*d*_ for which the *k*-scaled *i*_*c*_[*P*_*d*_ ← *P*_*o*_] *<* 0. **B**. Same construction as in A, but for *k* ≤ 1 (delay). Smaller *k* (0.75, 0.45, 0.25) pulls the bound back toward *P*_*o*_’s own favored vertex, requiring stronger evidence to commit. The committed regions at right (*k* = 0.75) are correspondingly smaller than in A, leaving a larger undecided region in the center of the simplex.

The flexibility of this approach is illustrated by including an “urgency” signal often used in decision models to account for a tendency to avoid long deliberation times found under some conditions [47]. The decision variable *P*_*d*_, like the bound *P*_*o*_, carries its own information compass. When referenced to the maximum-entropy belief *Q*, where the natural parameters vanish, that compass points along −*η*_*d*_, so the beliefs lying on it are those with *e*-coordinates *kη*_*d*_ for a scalar *k*. Rescaling the decision variable by *k* is therefore a move along its own compass, toward (or beyond) *Q* for *k <* 1 and away from it for *k >* 1 (Fig. 5). Accordingly, when *k* = 1, the decision variable is unchanged and commitment occurs at *P*_*o*_’s own fixed *i*_*c*_ = 0 line. When *k <* 1, the decision variable is pulled toward *Q*, so more evidence is required before commitment (delay). When *k >* 1, it is pushed away from *Q*, so less evidence suffices (urgency; 31). Thus, a simple scaling of the decision variable’s compass defines a trade-off between urgency and delay, illustrating more generally that any movement along the compass axis, whether by evidence or by rescaling, has an immediate interpretation in terms of commitment.

#### Information seeking

Decision rules that end evidence accumulation, like the bound in a DDM, typically optimize the trade-off between speed and accuracy, for example to maximize the rate of correct (rewarded) responses. More generally, however, information seeking may rely on other objectives that govern when enough has been learned about one domain to direct attention elsewhere [22].

Here we propose one such objective that can be defined using the information compass. Whereas Bayesian surprise quantifies how much an observation has already changed a belief [48] and classical accounts value an experiment by its expected information gain [49], we use the compass to define a prospective criterion, such that information is collected until reaching just enough certainty to be surprised if something changes. This definition differs from active-inference accounts that score a policy by its expected free energy [50], given that the criterion here is a threshold on the belief itself rather than on anticipated information gain.

We illustrate this idea in the context of a recent study of information-seeking behaviors [51]. Participants in that study tended to sample information from parts of the task environment (tables of card decks) about which they were most uncertain, thereby reducing their overall uncertainty. However, this strategy was apparent mainly when overall uncertainty was relatively low. In contrast, when overall uncertainty was relatively high they tended to use an opposite strategy and sampled from parts of the environment about which they were least uncertain. The crossover point between these two strategies thus marks a natural transition, between sampling what is already best known when overall uncertainty is high and sampling what is least known once overall uncertainty has fallen.

This crossover point corresponded, on average, to a belief that can be thought of as an informationgathering bound. For a source with two possible states, this bound (*P*_*b*_) can be derived directly from the compass as the belief at which expected *i*_*c*_ and expected Δ*φ* are equal, averaged over all possible destination beliefs, taken as uniformly distributed over the simplex (Fig. 6C). For two alternatives, this crossover occurs at *p* ≈ 0.68 for the favored alternative, an entropy of 0.63 nats.

**Fig. 6.**
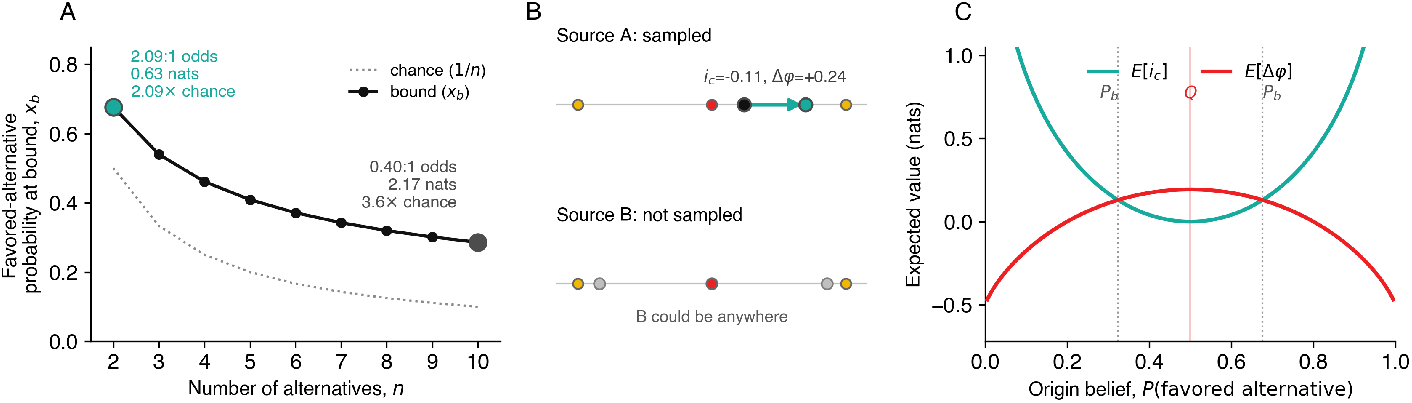
The information-gathering bound, *P*_*b*_. **A.** *P*_*b*_ (the favored-alternative probability at which expected *i*_*c*_ and expected Δ*φ* are equal) as a function of the number of alternatives a source can take, *n*, compared to chance (1*/n*, dotted). More alternatives require less absolute confidence, but a larger fold-improvement over chance, to cross the bound. **B**. When considering multiple, independent sources of information, *P*_*b*_ is a per-source property: an update to one source (A) has the same *i*_*c*_ and Δ*φ* regardless of a second, independent source’s (B) current state, because both quantities are additive across independent beliefs. **C**. Expected *i*_*c*_ (teal) and Δ*φ* (red) for a two-state (*n* = 2) source, crossing at *P*_*b*_ (dotted lines, symmetric about *Q*). Between these points, expected belief change is dominated by gains in certainty. Outside them, it is dominated by *i*_*c*_. Doubling the single-source bound (1.26 nats) closely matches the 1.28-nat switch reported by Abir and colleagues [51].

More generally, *P*_*b*_ depends on the number of alternatives a source can take. Sources with more possible states cross their bound at lower absolute confidence, but a larger fold-improvement over chance (Fig. 6A). *P*_*b*_ is also a strictly local, per-source property. Because both *i*_*c*_ and Δ*φ* are additive across independent beliefs, the bound for one source is unaffected by how many other, independent sources are being tracked at the same time (Fig. 6B). This additivity also lets us connect *P*_*b*_ quantitatively to Abir et al. [51]: their “overall uncertainty” statistic, which is the summed entropy of the two independent, binary beliefs (tables) shown on each trial, is by the same additivity exactly the entropy of the joint belief over both tables. If both tables are simultaneously at their own bound, the switch is predicted at 2 × 0.63 = 1.26 nats, closely matching their empirically estimated value of 1.28 nats; Fig. 6C.

Although the value of *P*_*b*_ itself depends on *n* (Fig. 6A), the criterion that defines it does not. Because *i*_*c*_ and Δ*φ* decompose the KL divergence between an origin and a destination belief exactly (Eq. 2), the crossover condition *E*[*i*_*c*_] = *E*[Δ*φ*] is equivalent to 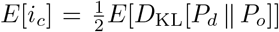. Thus, *P*_*b*_ is the belief at which, when averaging over possible new information, the compass-aligned component of belief change and the change in certainty contribute equally to expected surprise.

This interpretation suggests that the information-sampling strategies identified by Abir and colleagues [51] reflect a natural information-gathering bound. Between *Q* and *P*_*b*_, expected belief change is dominated by gains in certainty (Δ*φ*). There, most new information would make the belief more certain, but the belief is still too uncertain for that information to clearly support or contradict it. Beyond *P*_*b*_, expected belief change is instead dominated by its compass-aligned component (*i*_*c*_). There, new information bears mainly on whether the current belief is correct. Accordingly, people first seek information that builds certainty about what is already best known and, once past the bound, shift to discovering what the current belief fails to capture. Put another way, this bound reflects the idea that learning about what is not known first requires something to be known.

## Discussion

Probabilistic reasoning allows us to deal with the uncertainty inherent to our world by balancing deliberation (how and when we update our uncertain beliefs) with commitment (how and when we act on those beliefs). Most of what is known about these processes in the brain comes from binary choice tasks, in which the relevant belief reduces to a single scalar quantity (often a log-odds ratio) [5, 6]. Here we sought to generalize principles derived from those tasks to the richer, higher-dimensional beliefs that characterize real-world cognition.

We introduced the “information compass,” a construct grounded in information geometry [11, 12] that assigns to any categorical belief an axis in information space along which updates reinforce or repudiate that belief. By projecting potentially high-dimensional updates onto this single axis, the compass recovers the scalar log-odds of two-alternative tasks as a special case. We showed that this construct provides a common treatment of deliberation and commitment across decision-making, learning, and information-seeking problems. Here we discuss the advantages of using this geometric formulation, how its two coordinate systems might map onto known features of brain organization, and several open questions.

Treating beliefs as points on a manifold with dual coordinate systems, rather than working directly with whatever coordinates a given problem happens to be posed in, provides two important benefits. First, each coordinate system makes a particular kind of operation linear (mixture in *m*-coordinates, addition in *e*-coordinates). Performed in the coordinate system in which it is linear, an operation keeps the same form as the number of alternatives grows, affecting only the size of the vectors it acts on. For example, our two-step hazard-rate update, a convex combination in *m*-coordinates (Eq. 5) followed by a vector addition in *e*-coordinates (Eq. 6), extends with no new derivation to switching among any number of states (Methods M3). Likewise, the same sign-of-*i*_*c*_ criterion that defines a decision bound for two alternatives applies to any number of alternatives, without solving a new value function each time a state is added [31, 44]. These results are not merely convenient bookkeeping but instead reflect fundamental invariances of these geometric formulations as the number of alternatives increases.

Second, the same principle gives *i*_*c*_ a special status in information geometry. As a function of the destination belief’s *m*-coordinates, the KL divergence is nonlinear: it is the gap between the negentropy *φ*[*P*_*d*_] and the tangent line to *φ* at *P*_*o*_ (Methods M2). *i*_*c*_ is the linear part of this gap, set by the tangent line’s slope, *η*_*o*_, which depends only on *P*_*o*_. Δ*φ* is the nonlinear part, which contains all of the curvature. Accordingly, *i*_*c*_ can always be computed as a single inner product with −*η*_*o*_, regardless of how many alternatives are involved, even though the KL divergence it decomposes cannot generally be computed this way [11, 12]. This property makes *i*_*c*_ akin to natural gradients, mirror descent, and other approaches that exploit information geometry to make learning and inference efficient in high-dimensional spaces [52, 53]. These two coordinate systems may also align with different neural computations, suggesting an interesting, if speculative, mapping of these ideas onto functional brain organization. Operations native to *m*-coordinates, such as marginalizing out nuisance variables, mixing across competing hypotheses, or the *m*-step of our hazard-rate update, are weighted combinations of probabilities. This class of operation can be implemented by computations related to divisive normalization. Quadratic divisive-normalization circuits have been shown to implement marginalization of a probabilistically coded population under specific, biologically motivated coding assumptions [54]. These findings are consistent with normalization’s broader status as a canonical computation that is especially prominent in early sensory cortex [55].

In contrast, operations native to *e*-coordinates are naturally additive in log-probabilities. For example, the *e*-step of the hazard update is the addition of a new log-likelihood ratio to a running total. This form of update matches well-established findings that neurons in certain subcortical and cortical areas accumulate log-likelihood-ratio-like evidence toward a bound [8, 19, 56–58], in a manner closely related to the sequential probability ratio test [17]. Natural parameters have also been proposed more generally as a currency for representing uncertainty in population codes, independent of any decision context [59, 60]. This idea is consistent with the common use of logarithmic transformations in neural codes, from the log-polar map of primary visual cortex [61] to compressed codes for number [62, 63] and elapsed time [64, 65], and with evidence that perception follows the Fisher-information geometry of sensory codes [66, 67]. Whether these correspondences reflect only a superficial resemblance between our derivations and these known neural computations, or instead point to *m*- and *e*-coordinates as genuine organizing principles of brain function, is a question for further work.

Beyond these correspondences, the framework makes several predictions. First, because *i*_*c*_ is signed, neural and physiological signals associated with belief updating should distinguish updates that reinforce a belief from those that repudiate it, even when the two are matched for unsigned measures of surprise such as the KL divergence [13]. Second, in changing environments with more than two alternatives, signals related to expected volatility could track the hazard-dependent contribution to *i*_*c*_, which would repudiate the current belief, whereas signals related to new evidence would track the evidence-dependent contribution, which can take either sign [33].

We have restricted our treatment to discrete, categorical beliefs and to settings in which the reference belief *P*_*r*_ is fixed and externally supplied. The same dual-coordinate structure extends to continuous exponential-family distributions, suggesting that the same principles should carry over to richer classes of belief than we consider explicitly. However, extending the compass to other forms of structured beliefs, such as those with factorized or hierarchical dependencies among variables, is not automatic and would require specifying how the compass itself decomposes across that structure. Likewise, our motivated-behavior example combined two independently preferred variables into a single joint *P*_*r*_ by assumption. A fuller treatment would need to specify how *P*_*r*_ is itself acquired, updated, or arbitrated when multiple, potentially competing goals are present, rather than fixed in advance.

Our decomposition of belief change into a change in certainty (Δ*φ*) and a signed, compass-aligned component (*i*_*c*_) is reminiscent of the active-inference formalism, which decomposes expected free energy into epistemic (uncertainty resolution) and pragmatic (preference satisfaction) terms [3, 50]. However, neither term explicitly carries *i*_*c*_’s signed distinction between updates that reinforce and those that repudiate a given belief. Whether the two decompositions can be formally unified, for instance by giving expected free energy’s epistemic term a signed, *i*_*c*_-like refinement, and whether such a refinement would predict distinct behavioral signatures, such as a sharper transition between exploratory and exploitative phases of information seeking (Fig. 6), are open questions.

In summary, casting deliberation and commitment in the dual-coordinate language of information geometry does more than repackage familiar results in new notation. It exposes shared structure across problems that otherwise look unrelated; provides a single, computationally efficient toolkit that scales to any number of alternatives; and offers a speculative, but testable, link between this structure and established features of neural computation. A key question now is whether the information compass can retain these advantages in the higher-dimensional, naturalistic settings that characterize real-world probabilistic reasoning. If so, it could provide a common geometric language for understanding how neural populations represent, update, and act on complex probabilistic beliefs.

## Methods

### M1 Information geometry basics and notation

A key feature of information geometry is the use of dual coordinate systems, often referred to as *m*- (mixture or expectation-parameter) coordinates and *e*- (exponential or natural-parameter) coordinates, to represent any probability distribution [11, 12]. That is, any distribution *P* (here we focus on *n*-category (categorical) probability distributions over discrete states, of which the Bernoulli distribution is the *n* = 2 case) can be treated as a point characterized by expectation parameters *p* that determine its location in a probability simplex and by natural parameters *η* that determine its location in parameter space (Extended Data Fig. 1). The parameters *p* and *η* are connected by the Legendre transformations of information geometry [11, 12], as follows.

The *e*-coordinate representation *η* of *P* requires using one of the *n* states as a reference. Taking the last state (*s*_*n−*1_) as the reference yields:

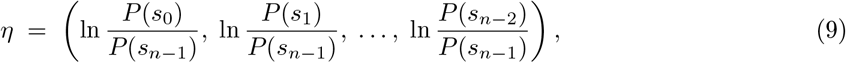

with dimension *n* − 1.

The corresponding *m*-coordinate representation *p* (expectation parameters) of *P* is:

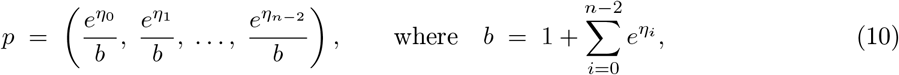

which simplifies to *p* = *P* (*s*_0_), *P* (*s*_1_), …, *P* (*s*_*n−*2_), with dimension *n* − 1.

The two functions allowing (Legendre) transformations between *η* and *p* are the negentropy function, *φ*, and the log-partition function, *ψ* (also known in the statistics literature as the cumulant generating function). The negentropy function can be written as:

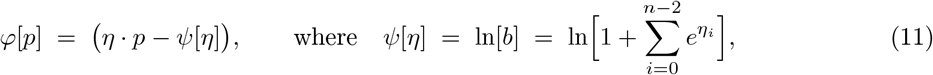

and *η* on the right-hand side denotes the natural-parameter representation of the same distribution as *p*. Substituting the expressions for *η, p*, and *ψ* shows that *φ* is the negative of the Shannon entropy *H*, summed over all *n* states: 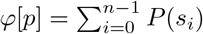 ln *P* (*s*_i_ ) = −*H*[*P* ]. Because *p* and *P* determine each other, we write *φ*[*P* ] and *φ*[*p*] interchangeably. Because *φ* and *ψ* are Legendre duals, *η* = ∇*φ*[*p*] and *p* = ∇*ψ*[*η*], where ∇ is the gradient operator. The special point *η* = 0 (equivalently, *p* uniform over all *n* states) is the maximum-entropy distribution, which we denote *Q*.

The *e*-geodesic through *Q* and a belief *P* (the compass axis; Fig. 1B) is the line *η*(*t*) = *t η*, where *η* is *P* ‘s *e*-coordinate representation. In *m*-coordinates, this line is

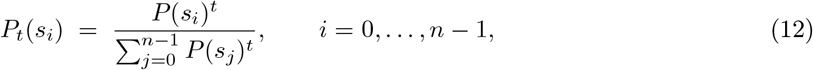

equivalently 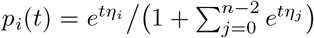 for *i* ≤ *n* − 2. As *t* → ∞, the belief concentrates on the state that *P* considers most probable or, if several states tie for most probable, spreads equally over just those states. As *t* → −∞, it concentrates in the same way on the least probable state(s). The endpoints of the compass axis are therefore vertices of the simplex only when the most (or least) probable state is unique.

Because *p* and *η* are Legendre-dual coordinates, the ordinary (Euclidean) inner product between a tangent vector at a given point expressed in *e*-coordinates and another tangent vector at the same point expressed in *m*-coordinates is exactly the pairing that recovers the Fisher-information metric of this statistical manifold (the natural, coordinate-invariant notion of distance on it) without ever needing to compute that metric, or its inverse, explicitly. This inner product is used to define *i*_*c*_ (Methods M2).

### M2 Derivation of *i*_*c*_ and its relationship to other information-theoretic quantities

This section derives *i*_*c*_ from first principles, as an inner product between two tangent vectors expressed in dual coordinates, and shows how it relates to the Kullback–Leibler (KL) divergence and to change in entropy (i.e., “certainty” gain).

The generalized Pythagorean theorem uses the duality of information geometry to define relationships between any three beliefs (e.g., *P*_*o*_, *P*_*d*_, and *P*_*r*_) in terms of the tangent vector from *P*_*o*_ to *P*_*r*_ in *e*-coordinates and the tangent vector from *P*_*o*_ to *P*_*d*_ in *m*-coordinates (Extended Data Fig. 2). Here we assume that *P*_*r*_ = *Q*, which is the maximum-uncertainty distribution and the only distribution whose *e*-coordinate representation is null (i.e., *η*_*Q*_ = 0), and define *i*_*c*_[*P*_*d*_ ← *P*_*o*_] as follows:

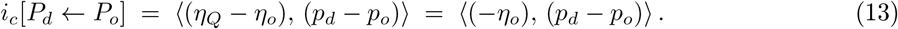

For a general reference belief, replacing *η*_*Q*_ with *η*_*r*_ gives the realigned compass *i*_*c,r*_[*P*_*d*_ ← *P*_*o*_] = ⟨(*η*_*r*_ − *η*_*o*_), (*p*_*d*_ − *p*_*o*_)⟩ of Eq. 8.

Note that this equation requires both *P*_*o*_ and *P*_*r*_ to assign nonzero probability to every state. A belief that has entirely ruled out even one state has no finite compass direction, because computing *η*_*o*_ takes the logarithm of that state’s probability. *P*_*d*_ carries no such restriction and may lie anywhere on the simplex, including its boundary.

**Extended Data Fig. 1.**
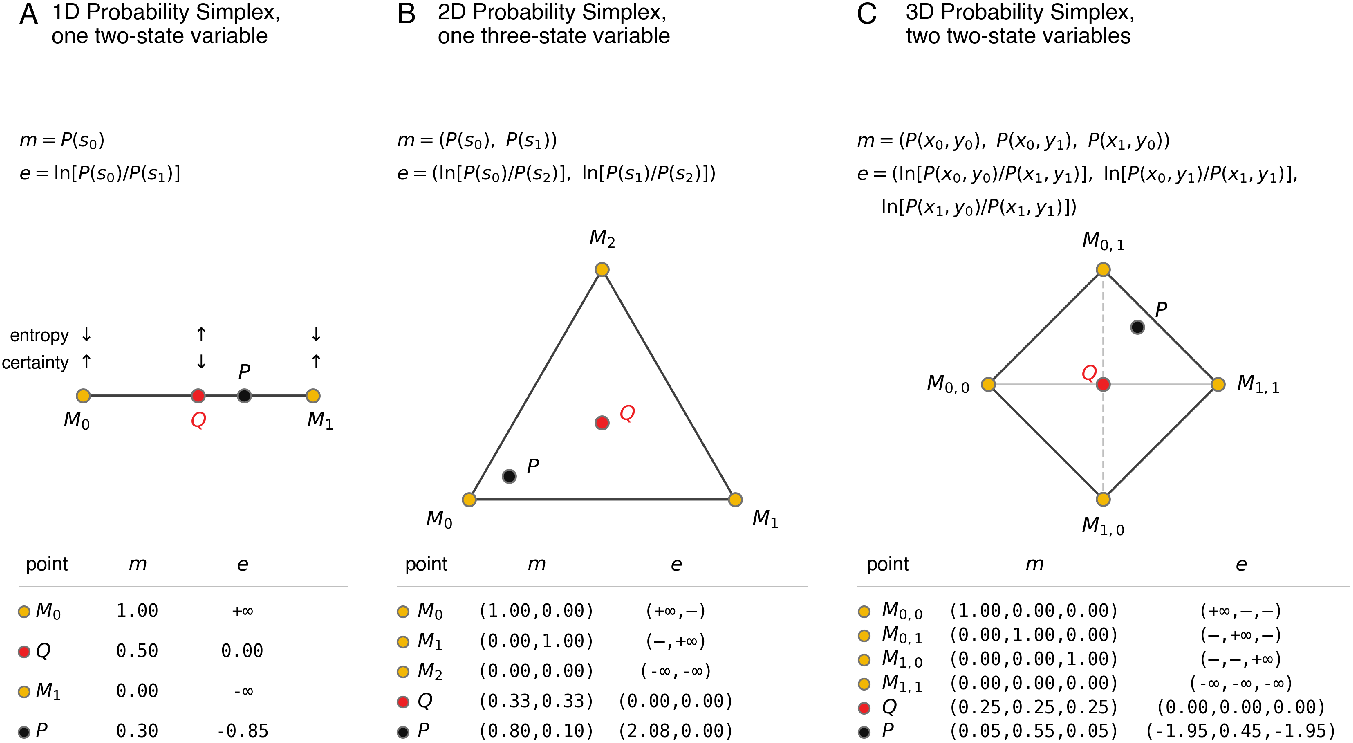
Categorical probability distributions represented in the simplex. An *n*-category distribution (e.g., *P* ) can be depicted as a point (*p*) in a simplex with *n* − 1 dimensions, each describing the expected probability (i.e., a value between zero and one) of one of the categories (the “expectation parameters”; note that once *n* − 1 probabilities have been specified, the full probability distribution is known, because all of the probabilities must sum to one). **A**. The simplex for a two-category belief is a line segment, consistent with the use of a scalar decision variable for modeling two-alternative tasks. **B**. The simplex for a three-category belief is a triangle (a two-dimensional region). **C**. The simplex for a four-category belief is a three-dimensional volume (a tetrahedron, here drawn in its symmetric form with the vertical dashed line representing an edge further away in depth than the horizontal solid line). In each panel, the red point represents the single maximum-entropy distribution (*Q*), yellow points represent all possible minimum-entropy distributions (*M*_*i*_), and the point labeled *P* represents an example probability distribution. Beneath each panel’s title, the exact reduced *m*- and *e*-coordinate expressions for that panel’s categories are given explicitly (Methods M1: *m* drops the last category and *e* is its log-ratio to that category, used as the *e*-coordinate reference). States are indexed from zero, as in the text, and the last state is the *e*-coordinate reference; in C, each of the four states is a pair (*x*_*i*_, *y*_*j*_ ) of values of two two-state variables, with (*x*_1_, *y*_1_) as the reference. The table below each panel gives the resulting *m*- and *e*-coordinates of every labeled point. *M*_*i*_ points are minimum-entropy vertices, where *e*-coordinates diverge to ±∞; entries marked “–” are additionally indeterminate (0*/*0) because more than one category vanishes together at that vertex.

To relate Eq. 13 to familiar information-theoretic quantities, we first write it as a sum over all *n* states. Each component of *η*_*o*_ is ln *P*_*o*_(*s*_*i*_) − ln *P*_*o*_(*s*_*n−*1_), and because both beliefs sum to one, 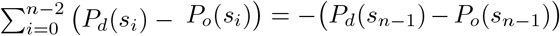. The terms involving the reference state *s*_*n−*1_ therefore fold back into the sum as its final (*i* = *n* − 1) term, giving:

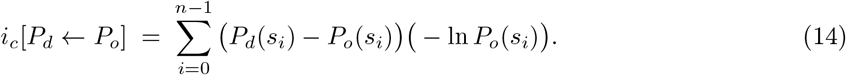

That is, *i*_*c*_ is the change in the probability assigned to each state, weighted by that state’s surprise under the origin belief. Because the changes in probability sum to zero, subtracting the origin belief’s average surprise (its entropy, *H*[*P*_*o*_] = −*φ*[*P*_*o*_]) from every weight leaves this sum unchanged. Thus, *i*_*c*_ *>* 0 when probability shifts toward states that are more surprising than average under *P*_*o*_, and *i*_*c*_ *<* 0 when it shifts toward states that are less surprising than average.

Distributing the product splits Eq. 14 into two familiar terms: 1) − ∑_*i*_ *P*_*d*_(*s*_*i*_) ln *P*_*o*_(*s*_*i*_), which is the cross-entropy *H*[*P*_*d*_, *P*_*o*_]; and 2) ∑_*i*_ *P*_*o*_(*s*_*i*_) ln *P*_*o*_(*s*_*i*_), which is the negentropy *φ*[*P*_*o*_] (Methods M1). Adding and subtracting *φ*[*P*_*d*_] = ∑_*i*_ *P*_*d*_(*s*_*i*_) ln *P*_*d*_(*s*_*i*_) then gives:

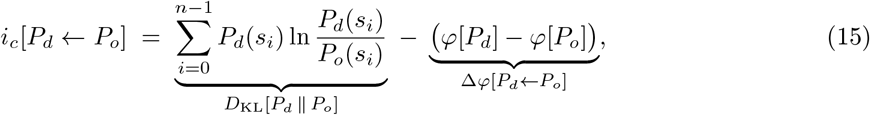

where *D*_KL_[· ∥ ·] is the Kullback–Leibler divergence and Δ*φ*[*P*_*d*_ ← *P*_*o*_] (written Δ*φ* for short below, because *P*_*o*_ and *P*_*d*_ are fixed throughout this section) is the gain in certainty (the reduction in entropy) from *P*_*o*_ to *P*_*d*_. Eq. 15 is Eq. 2 of the Results. Equivalently, *D*_KL_[*P*_*d*_ ∥ *P*_*o*_] = *H*[*P*_*d*_, *P*_*o*_] − *H*[*P*_*d*_] and *i*_*c*_ = *H*[*P*_*d*_, *P*_*o*_] − *H*[*P*_*o*_]: both compare the same cross-entropy, but the KL divergence compares it to the destination belief’s entropy, whereas *i*_*c*_ compares it to the origin belief’s entropy. Moving Δ*φ* to the other side expresses the KL divergence as the sum of the two components of the update:

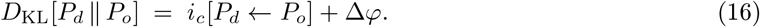

In addition, *D*_KL_[*P*_*d*_ ∥ *P*_*o*_], expressed as a function of the *m*-coordinates *p*_*d*_ and *p*_*o*_, is exactly the Bregman divergence generated by the negentropy potential *φ* [11, 12]: *D*_KL_[*P*_*d*_ ∥ *P*_*o*_] = *φ*[*P*_*d*_] − *φ*[*P*_*o*_] − ⟨*η*_*o*_, (*p*_*d*_ − *p*_*o*_)⟩, using ∇*φ*[*P*_*o*_] = *η*_*o*_. Geometrically, this divergence is the gap between *φ*[*P*_*d*_] and the tangent line (for *n >* 2, the tangent hyperplane) to *φ* at *P*_*o*_, evaluated at *p*_*d*_. This construction identifies *i*_*c*_ as the linear term of a standard Bregman decomposition, situating it within the broader family of natural-gradient and mirror-descent methods used elsewhere in statistics and machine learning for efficient inference and learning in high-dimensional spaces.

Eq. 16 can also be written entirely in terms of KL divergences, because Δ*φ* measures how far each belief is from *Q*. Because *Q* is uniform over the *n* states, *D*_KL_[*P* ∥ *Q*] = Σ_*i*_ *P* (*s*_*i*_) ln (*nP* (*s*_*i*_)) = ln(*n*) + Σ_*i*_ *P* (*s*_*i*_) ln *P* (*s*_*i*_) = ln(*n*) + *φ*[*P* ] for any belief *P*, using Σ_*i*_ *P* (*s*_*i*_) = 1. The constant ln(*n*) cancels in the difference between two beliefs, so:

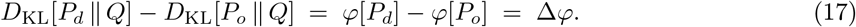

Substituting Eq. 17 for Δ*φ* in Eq. 16 and solving for *D*_KL_[*P*_*d*_ ∥ *Q*] gives a form analogous to the law of cosines:

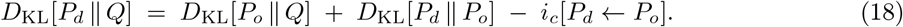

When *i*_*c*_ = 0, the update is orthogonal to the compass axis and Eq. 18 reduces to the Pythagorean theorem of information geometry [11]. Thus, nonzero values of *i*_*c*_ measure the departure from orthogonality. Repeating the same steps with a general reference belief *P*_*r*_ in place of *Q* gives *i*_*c,r*_[*P*_*d*_ ← *P*_*o*_] = ⟨(*η*_*r*_ − *η*_*o*_), (*p*_*d*_ − *p*_*o*_)⟩ = *D*_KL_[*P*_*d*_ ∥ *P*_*o*_] + *D*_KL_[*P*_*o*_ ∥ *P*_*r*_] − *D*_KL_[*P*_*d*_ ∥ *P*_*r*_], which is the realigned compass of Eq. 8.

For two-state beliefs, this decomposition also has a direct graphical reading on the negentropy curve *φ* and its tangent line at *P*_*o*_ (Extended Data Fig. 2A). The tangent line, *T* (*x*) = *φ*[*P*_*o*_] + *η*_*o*_(*x* − *p*_*o*_), extended to *P*_*d*_’s *m*-coordinate *p*_*d*_, gives the negative cross-entropy, *T* (*p*_*d*_) = −*H*[*P*_*d*_, *P*_*o*_]. The vertical gap between this tangent line and the curve at *p*_*d*_ is *D*_KL_[*P*_*d*_ ∥ *P*_*o*_] = *φ*[*P*_*d*_] −*T* (*p*_*d*_), which is never negative because *φ* is convex. A horizontal line at *P*_*o*_’s own height, *φ*[*P*_*o*_], splits this gap into two signed pieces: *φ*[*P*_*o*_] −*T* (*p*_*d*_), measured from the tangent line to the horizontal line, equals *i*_*c*_[*P*_*d*_ ← *P*_*o*_]; and *φ*[*P*_*d*_] − *φ*[*P*_*o*_], measured from the horizontal line to the curve, equals Δ*φ*[*P*_*d*_ ← *P*_*o*_]. Each piece is positive when it points upward. When both are positive, the horizontal line falls within the gap and the two pieces stack to form it. When one is negative, the horizontal line falls outside the gap. For example, the *P*_*d*_ shown in Extended Data Fig. 2A lies on the far side of *Q* from *P*_*o*_ but is less certain than *P*_*o*_, so *φ*[*P*_*d*_] *< φ*[*P*_*o*_], Δ*φ <* 0, and *i*_*c*_ exceeds *D*_KL_[*P*_*d*_ ∥ *P*_*o*_].

### M3 The two-step (*m*-then-*e*) derivation of the hazard update

#### Switching among *n* states: the exact Bayesian forward step

For a belief over *n* discrete states, let *q*(*s*_*i,t*_) ≡ *p*(*s*_*i*_ | *x*_1:*t*_) denote the posterior probability of state *s*_*i*_ given all evidence collected through time *t*, and assume a probability *h* that the state switches at each time step, redistributed uniformly over the other *n* − 1 states. Exactly as for the two-state case [33], Bayes’ rule together with the sum and product rules of probability give the forward recursion:

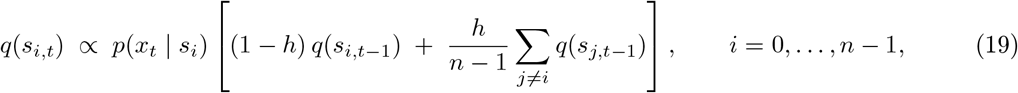

which is the forward step of the Baum–Welch algorithm for a Hidden Markov Model with *n* states and this transition kernel [70]. Because Σ_*j ≠ i*_ *q*(*s*_*j,t−*1_) = 1 − *q*(*s*_*i,t−*1_), the bracketed term is exactly *P*_*h*_(*s*_*i*_ ) from Eq. 5; taking logarithms recovers the (*e*-coordinate) evidence term exactly as in the two-state derivation. At *n* = 2, the sum over *j* ≠ *i* has only one term, so Eq. 19 reduces exactly to Eq. 4.

**Extended Data Fig. 2.**
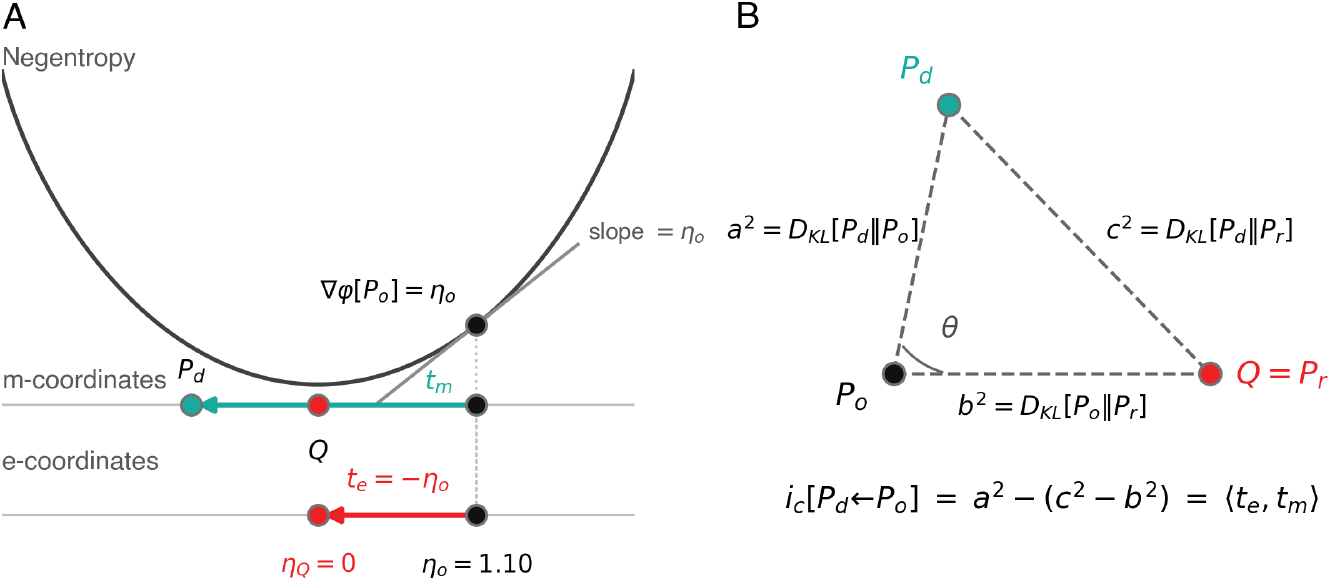
Derivation and properties of the information compass. **A**. Geometric interpretation of the information compass for a two-state belief. Top: the negentropy function *φ* over the 1D probability simplex (*m*-coordinates), with an origin belief *P*_*o*_ (black), a destination belief *P*_*d*_ (teal; these distributions might correspond to a Bayesian prior and posterior, respectively, but here we use more general terms to emphasize that the update does not need to be based on Bayesian principles), and the maximum-entropy belief *Q* (red), located at the trough of *φ* and here used as the reference belief (*P*_*r*_ ). The gradient (∇) of *φ* at *P*_*o*_ with respect to the *m*-coordinates equals its natural-parameter coordinate, ∇*φ*[*P*_*o*_] = *η*_*o*_ (Methods M1); *t*_*m*_ is the tangent vector of the *m*-geodesic from *P*_*o*_ to *P*_*d*_. Bottom: the same beliefs in natural-parameter space (*e*-coordinates), plotted at the same horizontal position as their *m*-coordinate above so that a belief’s *m*- and *e*-coordinates align vertically; *t*_*e*_ = −*η*_*o*_ is the tangent vector of the *e*-geodesic from *P*_*o*_ to *P*_*r*_ = *Q* (*η*_*Q*_ = 0). **B**. The generalized Pythagorean theorem in information geometry expresses the KL divergence as the squared distance between vertices of a schematic triangle defined by three probability distributions (*P*_*o*_, *P*_*d*_, and *P*_*r*_ = *Q*; note that this drawing is a schematic of the KL-divergence relationships between the three beliefs and not a literal embedding, given that KL divergence is not a Euclidean distance). *i*_*c*_ follows directly from this relationship as *a*^2^ − (*c*^2^ − *b*^2^), which is exactly the inner product ⟨*t*_*e*_, *t*_*m*_⟩ between the two tangent vectors defined in A. As such, *i*_*c*_ is the signed, unnormalized projection (inner product) of the update onto the compass, related to but not equal to the cosine of the angle *θ* at the origin-belief vertex (*P*_*o*_), since no norm has been chosen for either coordinate system; each geodesic is straight only in its own affine coordinates, and the two are linked by the duality theorem of information geometry [10–12, 68, 69] (full algebraic derivation in Methods M2). See Fig. 2B for concrete examples of *i*_*c*_ *<* 0, *i*_*c*_ ≈ 0, and *i*_*c*_ *>* 0 on a probability simplex.

**Extended Data Fig. 3.**
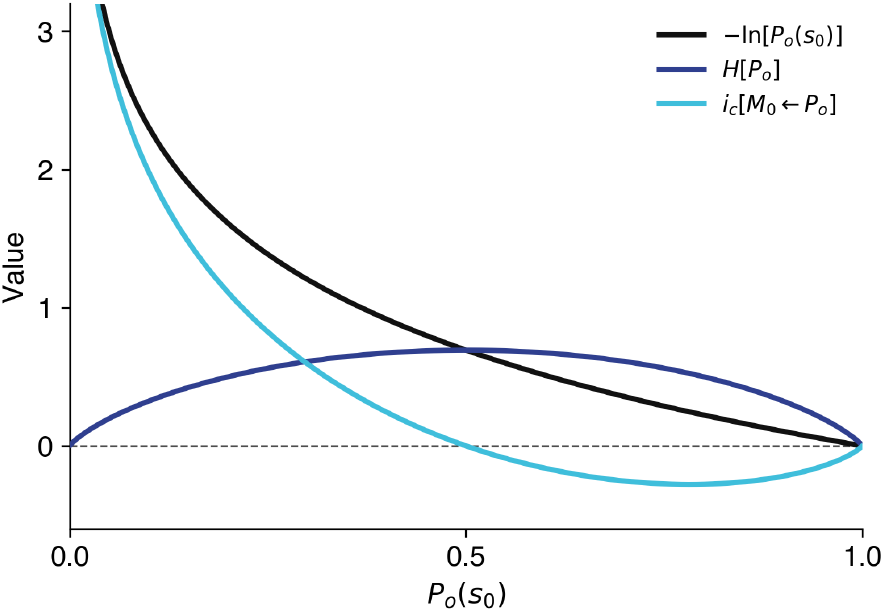
Surprise, entropy, and *i*_*c*_. Setup: a belief *P*_*o*_ about two states, with the two minimum-entropy (maximum-certainty) beliefs *M*_0_ = {*s*_0_ = 1, *s*_1_ = 0} and *M*_1_ = {*s*_0_ = 0, *s*_1_ = 1} at the simplex vertices, *Q* at the trough (maximum entropy, used as the reference belief *P*_*r*_), and the destination fixed at *P*_*d*_ = *M*_0_. Three quantities are plotted against *P*_*o*_(*s*_0_) for this fixed destination: 1) surprise, −ln[*P*_*o*_(*s*_0_)] = *D*_KL_[*M*_0_ ∥ *P*_*o*_] (black); 2) entropy, *H*[*P*_*o*_] (dark blue); and 3) *i*_*c*_[*M*_0_ ← *P*_*o*_] (light blue). These quantities satisfy *D*_KL_[*M*_0_ ∥ *P*_*o*_] = *i*_*c*_[*M*_0_ ← *P*_*o*_] + Δ*φ*[*M*_0_ ← *P*_*o*_] (Eq. 2), with Δ*φ*[*M*_0_ ← *P*_*o*_] = *φ*[*M*_0_] − *φ*[*P*_*o*_] = −*φ*[*P*_*o*_] = *H*[*P*_*o*_], because *φ*[*M*_0_] = 0 exactly at a vertex. Thus, the gap between surprise and *i*_*c*_ here is always exactly *H*[*P*_*o*_], and this gap does not depend on which vertex (*M*_0_ or *M*_1_) is chosen as the destination, because *φ* is 0 at either. In contrast, *i*_*c*_ varies with *P*_*o*_(*s*_0_) and is signed: a negative *i*_*c*_ indicates that, on average, more probable states are becoming even more probable, which is not captured by surprise.

This transition kernel assumes that every state exits at the same total rate *h*, split evenly across the other *n* − 1 states. More generally, a fully general transition-rate matrix includes a separate rate *ϵ*_*ij*_ governing switches from state *j* to state *i* for every ordered pair *i*≠ *j* [34]. The uniform-*h* kernel used here is the symmetric, exchangeable special case of that matrix (*ϵ*_*ij*_ = *h/*(*n* − 1) for all *i ≠ j*), and this symmetry is what makes *P*_*h*_ a mixture of exactly two fixed points, *P*_*o*_ and 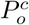 (Eq. 5). This mixture is thus a one-parameter, one-dimensional family, corresponding to an *m*-geodesic. A general *ϵ*_*ij*_ matrix has no single such target: because arbitrary switching rates do not necessarily privilege any one direction away from *P*_*o*_, the resulting update is a linear map with as many free parameters as there are ordered state pairs and is not confined to a one-dimensional family in either coordinate system. Note that the additive decomposition of *i*_*c*_ below does not depend on this symmetry (see below).

Taking *s*_*n−*1_ as the reference state and writing *p*_*i*_ ≡ *P*_*o*_(*s*_*i*_), Eq. 5 can be written entirely in terms of *p*_*i*_ and a single contraction factor *c* ≡ 1 − *hn/*(*n* − 1):

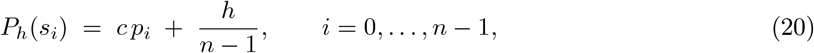

so that *P*_*h*_ − *P*_*o*_ = (1 − *c*)(*Q* −*P*_*o*_). Thus, the entire update moves along the straight line from *P*_*o*_ through *Q*, reaching *Q* when *c* = 0; i.e., at *h* = (*n* − 1)*/n*, which is = 0.5 when *n* = 2 and approaches 1 as *n* increases.

Converting Eq. 20 to *e*-coordinates gives

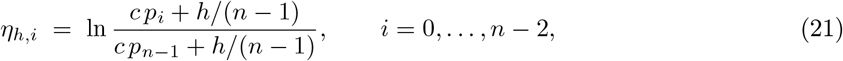

which reduces exactly to Eq. 2 of ref. [33] at *n* = 2 (where *c* = 1 −2*h*) and inherits both of its defining regimes for any *n*. Specifically, near *Q* (*p*_*i*_ ≈ 1*/n*), *η*_*h,i*_ ≈ *c η*_*o,i*_, representing a leaky-accumulator regime with leak 1 − *c* (see Eq. 3a of ref. [33]). As *P*_*o*_ approaches the vertex favoring *s*_*i*_ (*p*_*i*_ → 1), *η*_*h,i*_ approaches ln[(*n* −1)(1 − *h*)*/h*] regardless of how extreme *P*_*o*_ already was, representing a stabilizing boundary (see Eq. 3b,c of ref. [33]).

#### Additive decomposition of *i*_*c*_ (Eq. 7)

The two-step update raises a bookkeeping problem. The *m*-step’s effect is naturally a change in probability, whereas the *e*-step’s effect is a change in log-odds. Thus, comparing their relative contributions to the belief update requires converting both into one common scale. This conversion can be accomplished using *i*_*c*_, which quantifies any update as a signed displacement along one chosen axis. Once every step of an update is expressed this way, the resulting numbers can be added directly.

Because *i*_*c*_[*P*_*d*_ ← *P*_*o*_] = (−*η*_*o*_) · (*p*_*d*_ −*p*_*o*_) (Eq. 1) is linear in the destination belief’s *m*-coordinates for any fixed origin *P*_*o*_, it can be applied separately to the *m*-step’s displacement, Δ_*m*_ ≡ *p*_*h*_ − *p*_*o*_, and the *e*-step’s displacement, Δ_*e*_ ≡ *p*_*d*_ − *p*_*h*_, using the same −*η*_*o*_ throughout. These two displacements sum to the update’s net displacement regardless of the path’s geometry in between:

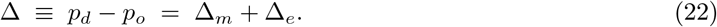

Applying *i*_*c*_ to each displacement with the same fixed −*η*_*o*_ gives *i*_*c*_[*P*_*h*_ ← *P*_*o*_] = (−*η*_*o*_) · Δ_*m*_ for the *m*-step and (−*η*_*o*_) · Δ_*e*_ for the *e*-step, and *i*_*c*_[*P*_*d*_ ← *P*_*o*_] = (−*η*_*o*_) · Δ for the full update. Because −*η*_*o*_ is the same vector throughout, and Δ = Δ_*m*_ +Δ_*e*_ (Eq. 22), the two steps sum to the same answer as the full update:

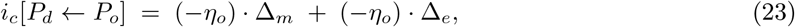

which is Eq. 7 (see Fig. 3C).

This derivation uses only linearity of the inner product and holds for any fixed reference belief chosen before the update begins. It also holds regardless of what generated *P*_*h*_; e.g., nothing in the argument requires Δ_*m*_ to come from a two-point mixture. The decomposition therefore applies unchanged to the general *ϵ*_*ij*_ switching matrix. That is, *i*_*c*_[*P*_*h*_ ← *P*_*o*_] = (−*η*_*o*_) · (*p*_*h*_ −*p*_*o*_) still isolates the *m*-step’s contribution on the same fixed axis as the *e*-step, and the two still sum exactly to *i*_*c*_[*P*_*d*_ ← *P*_*o*_], whether or not the first step traces a geodesic. The uniform-*h* case is what makes that first contribution expressible as a single mixture with a fixed target, but the compass’s bookkeeping does not require it.

#### Sign of the hazard-step contribution

When the reference is reset to the current belief at every timestep, as above, the *m*-step’s contribution has a fixed sign. Because *Q* is the global minimum of the negentropy potential *φ* (Methods M2), *i*_*c*_[*Q* ← *P*_*o*_] = *D*_KL_[*Q* ∥ *P*_*o*_] + (*φ*[*P*_*o*_] − *φ*[*Q*]) ≥ 0 for any *P*_*o*_, because both terms are non-negative. Given that 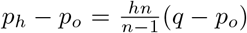 (Eq. 20) is a non-negative multiple of *q* − *p*_*o*_, the *m*-step’s contribution is 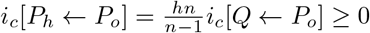. Hazard-driven mixing can therefore only ever repudiate the current belief, never reinforce it, regardless of *n*. This property depends not on the uniform kernel itself but on switching that does not favor particular states. For any doubly stochastic transition matrix (i.e., one under which, if all states start out equally likely, they remain equally likely), mixing cannot decrease entropy, so *i*_*c*_[*P*_*h*_ ←*P*_*o*_] = −*H*[*P*_*h*_, *P*_*o*_] − *H*[*P*_*o*_] ≥*H*[*P*_*h*_] − *H*[*P*_*o*_] ≥ 0 (Methods M2). This condition is also necessary: if switching makes some states more likely than others even from complete uncertainty, then there are origin beliefs near *Q* for which *i*_*c*_[*P*_*h*_ ← *P*_*o*_] *<* 0. Under such switching, only the evidence step can move *i*_*c*_ negative. Note also that for sufficiently high hazard rates (*h >* (*n* − 1)*/n* in the uniform case), the *m*-step alone carries the belief past *Q*, so a change of mind need not involve any evidence.

#### Local versus cumulative reference points

The decomposition above resets *P*_*o*_ at the start of every timestep, so *i*_*c*_[*P*_*d*_ ← *P*_*o*_] reports whether that one timestep’s update, on its own, reinforced or repudiated whatever was believed immediately beforehand. However, nothing in the additive argument above requires the reference to reset every timestep. Fixing *P*_*o*_ instead at a trial’s starting belief, *P*_*o*,0_, and never updating it, allows every subsequent mixing and evidence step, across an entire trial, to be scored against the same −*η*_*o*,0_. These values can be summed into one running total:

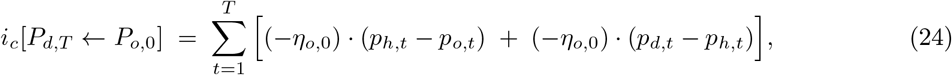

where *P*_*o,t*_ ≡ *P*_*d,t−*1_ is the belief entering timestep *t*. This cumulative reading answers a different question than the local one above: not whether the most recent update alone was confirming or disconfirming, but whether the process as a whole, evidence and volatility together, has moved the belief toward or away from the state(s) originally favored at trial onset. With this fixed reference, the sign guarantee for the *m*-step no longer holds: if the belief has come to favor a state that *P*_*o*,0_ disfavored, mixing toward *Q* can move it back toward *P*_*o*,0_, which counts as reinforcement (*i*_*c*_ *<* 0) on the original axis.

#### A compass toward a belief’s opposite

Consider instead a hypothetical change not in the state of the world but in one’s own belief: with probability *h*, the belief is replaced by its exact reversal. We define for any belief *P*_*o*_ its “opposite” 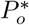, which together form a pair that, when combined, cancel each other out exactly to result in the maximum-entropy belief *Q*. This definition is straightforward in *e*-coordinates: the opposite of *η*_*o*_ (the *e*-coordinate representation of *P*_*o*_) is *η*^*\**^ = −*η*, such that *η*_*o*_ +*η*^*\**^ = 0 = *Q*; equivalently, 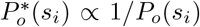. By this construction, *P*_*o*_, *Q*, and 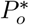 always lie along the same *e*-geodesic, for any *n*. Note that 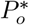 is generally distinct from 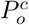 (Eq. 5): 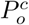 lies opposite *P*_*o*_ through *Q* on a straight line in *m*-coordinates, at 1*/*(*n* − 1) of *P*_*o*_’s distance from *Q*, whereas 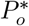 is its mirror image through *Q* in *e*-coordinates (straight in log-odds). The two coincide only at *n* = 2, where a single axis admits just one notion of ‘opposite’. For *n >* 2, the two definitions diverge.

Under these conditions, mixing *P*_*o*_ with 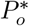 yields:

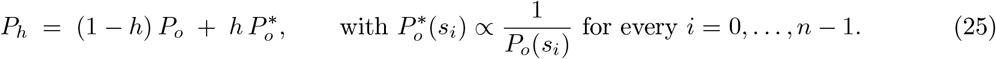

Here *P*_*h*_ is not linear in *P*_*o*_ for *n >* 2: mixing two beliefs and then taking the opposite of the result does not generally equal mixing their two, separately computed, opposites. Put another way, for *n >* 2 the opposite 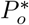 is a nonlinear function of *P*_*o*_, whereas at *n* = 2 it simply swaps the two probabilities. Thus, under these conditions Eq. 25 is best read not as an update rule meant to be applied repeatedly, but as a one-time blend between a belief and a single, hypothetical, complete reversal of it: *h* = 0 leaves *P*_*o*_ unchanged, *h* = 1 reaches the exact reversal 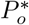, and intermediate *h* interpolate between them.

Because *P*_*o*_, *Q*, and 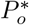 share one *e*-geodesic for any *n*, 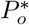 is a natural choice of reference belief (*P*_*r*_) for Eq. 8, giving a fixed reference direction that depends only on *P*_*o*_ itself:

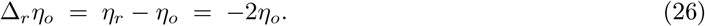

Because Δ_*r*_*η*_*o*_ = 2(−*η*_*o*_), the resulting *i*_*c*_ is exactly twice the *Q*-referenced *i*_*c*_ of Eq. 1: the axis and sign are unchanged, and only the interpretation of the endpoint differs. By the same linearity argument as above, this reference direction decomposes any two-step update additively, exactly as in Eq. 7, now interpreted as progress toward (or away from) a complete reversal of *P*_*o*_ rather than toward maximum uncertainty:

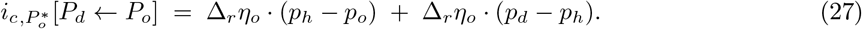

Positive values indicate that the update moved the belief toward having fully abandoned *P*_*o*_’s favored state for its exact opposite. Negative values indicate the reverse. This reading is available for any belief and any update, regardless of what generated *P*_*h*_ (e.g., via Eq. 5).

## Acknowledgements

We thank Pratik Chaudhari, Amrut Nadgir, Marc Miskin, and Eugenio Piasini for helpful feedback.

## Declarations

### Data availability

This is a theoretical study, so no new experimental data were generated.

### Code availability

Code used to generate the figures is available at https://github.com/TheGoldLab/ms_2026_bosulu_etal.

### Author contributions

J.B. conceived the framework and developed the formalism; J.B. and J.I.G. produced the figures; all authors wrote, reviewed, and edited the manuscript.

